# Transgenerational effect of mutants in the RNA-directed DNA methylation pathway on the triploid block

**DOI:** 10.1101/2020.09.08.288001

**Authors:** Zhenxing Wang, Nicolas Butel, Juan Santos-González, Lauriane Simon, Cecilia Wärdig, Claudia Köhler

## Abstract

Hybridization of plants that differ in number of chromosome sets (ploidy) frequently causes endosperm failure and seed arrest, a phenomenon referred to as triploid block. Mutation in *NRPD1*, encoding the largest subunit of the plant-specific RNA Polymerase IV (Pol IV), can suppress the triploid block. Pol IV generates short RNAs required to guide *de novo* methylation in the RNA-directed DNA methylation (RdDM) pathway. In this study, we found that the ability of mutants in the RdDM pathway to suppress the triploid block depends on their degree of inbreeding. While *nrpd1* is able to suppress in the first homozygous generation, mutants in *RDR2*, *NRPE1*, and *DRM2* require at least one additional round of inbreeding to exert a suppressive effect. Inbreeding of *nrpd1* was connected with a transgenerational loss of non-CG DNA methylation on sites jointly regulated by CHROMOMETHYLASES 2 and 3. Our data thus reveal that loss of RdDM function differs in its effect in early and late generations and that Pol IV acts at an early stage of triploid block establishment.

**One-sentence summary:** Inbreeding of mutants impaired in RdDM components transgenerationally enhanced their ability to suppress the triploid block.

## Introduction

Hybridization of plants that differ in ploidy frequently leads to seed arrest, a phenomenon referred to as the triploid block (Marks, 1966; Köhler et al., 2010). The triploid block is stablished in the endosperm, a nourishing tissue supporting embryo growth (Brink and Cooper, 1947; Johnston et al., 1980; Lin, 1984). The endosperm is typically a triploid tissue, derived after fertilization of the diploid central cell by one of the sperm cells (Li and Berger, 2012). In most flowering plant species, the endosperm initially develops as a syncytium and undergoes cellularization after a defined number of nuclear divisions (Costa et al., 2004). In *Arabidopsis thaliana*, as in many other flowering plant species, hybridizations of maternal plants with higher ploidy pollen donors causes failure of the endosperm to cellularize, leading to embryo arrest (Scott et al., 1998; Hehenberger et al., 2012). Sensitivity of the endosperm to parental genome dosage is closely connected to genomic imprinting, an epigenetic phenomenon resulting in the parental-specific expression of specific genes (Gehring et al., 2017; Batista and Kohler, 2020). Specifically, loss of function of the imprinted paternally expressed genes (PEGs) *ADMETOS*, *SUVH7*, *SUVH9*, *AHL10*, *PEG2*, *PEG9, PICKLE RELATED2* (*PKR2)*, and *PHERES1* (*PHE1*) are sufficient to suppress the triploid block (Kradolfer et al., 2013; Wolff et al., 2015; Huang et al., 2017; Inoue et al., 2017; Wang et al., 2018; Batista et al., 2019b), suggesting a causal role of imprinted genes in establishing the triploid block. Similarly, loss of function of the paternally-biased *NRPD1* gene, encoding the largest subunit of the plant specific RNA Polymerase IV (Pol IV), leads to suppression of the triploid block (Erdmann et al., 2017; Martinez et al., 2018). Pol IV is a central component of the RNA-directed DNA methylation (RdDM) pathway that establishes DNA methylation in all sequence contexts and maintains CHH methylation (H corresponds to A, C, or T) preferentially on small eukaryotic TEs (Matzke and Mosher, 2014; Zhang et al., 2018). Pol IV forms short transcripts of 26-45-nt in size that are converted into double stranded RNA by the RNA-DEPENDENT RNA POLYMERASE 2 (RDR2) (Blevins et al., 2015; Zhai et al., 2015). Double-stranded RNAs are then targeted by different DICER-LIKE (DCL) proteins to generate small RNAs (sRNAs) in the size range of 21-24-nt that are incorporated into ARGONAUTE (AGO) proteins. These sRNA-AGO complexes pair with Pol V-derived scaffold transcripts and recruit the DOMAINS REARRANGED METHYLTRANSFERASE2 (DRM2), which methylates DNA in all sequence contexts (Law and Jacobsen, 2010; Zhang et al., 2018; Panda et al., 2020; Wang et al., 2020). Pol IV is recruited to heterochromatic regions by SAWADEE HOMEODOMAIN HOMOLOGUE 1 (SHH1), which recognizes dimethylated histone H3 lysine 9 (H3K9me2) (Law et al., 2013; Zhang et al., 2013). Methylation on CHH positions can also be mediated by CHROMOMETHYLASE 2 (CMT2), which acts in a feedback loop with H3K9me2 (Zemach et al., 2013; Stroud et al., 2014). CMT2 can also target CHG positions, but at reduced efficiency (Stroud et al., 2014). The main CHG methyltransferase is CHROMOMETHYLASE 3 (CMT3), which like CMT2 maintains CHG methylation in a feedback loop with H3K9me2 (Lindroth et al., 2001; Jackson et al., 2002; Malagnac et al., 2002; Du et al., 2012; Zemach et al., 2013). Both, CMT2 and CMT3 preferentially target heterochromatic TEs, while the RdDM pathway preferentially targets short euchromatic TEs (Zemach et al., 2013). Maintenance of CG methylation requires METHYLTRANSFERASE 1 (MET1), which recognizes hemi-methylated symmetrical CG nucleotides (Law and Jacobsen, 2010; Zhang et al., 2018). Loss of paternal MET1 function suppresses the triploid block (Schatlowski et al., 2014), similar to aforementioned mutants in PEGs. Also the triple *suvh4/5/6* mutant that is deficient in the H3K9me2 methyltransferases KRYPTONITE (KYP, or SUVH4) and the redundantly acting SUVH5 and SUVH6, is a strong suppressor of the triploid block (Jiang et al., 2017). These data point that there is a connection between DNA methylation and the triploid block, but the precise mechanisms and targets remain to be identified.

Recent work suggests that suppression of the triploid block by mutants in the RdDM components *RDR2*, *DCL3*, *NRPE1*, and *DRM2* differs, depending on whether the diploid pollen (2n) is derived from tetraploid (4x) plants or from the *osd1* (*omission in second division 1*) mutant. Loss of *OSD1* causes an omission of the second meiotic division, leading to 2n pollen formation (d’Erfurth et al., 2009). While 4x mutants in *NRPE1*, *RDR2*, *DCL3*, and *DRM2* suppress the triploid block (Satyaki and Gehring, 2019), no suppressive effect of those mutants was found in the *osd1* background (Martinez et al., 2018). However, mutants in the Pol IV component *NRPD1* could similarly suppress the triploid block in 4x and *osd1* background (Martinez et al., 2018; Satyaki and Gehring, 2019), suggesting that there is a difference in the response to loss of RdDM function in *osd1* and 4x plants. Tetraploid RdDM mutants were generated from inbred mutants using colchicine treatment (Satyaki and Gehring, 2019), while RdDM mutants in *osd1* background were selected from segregating F2 populations (Martinez et al., 2018). Previous work in maize revealed that loss of Pol IV function causes a progressively enhanced loss of silencing over generations (Erhard et al., 2013), suggesting that it could make a difference whether using first generation homozygous RdDM mutants or highly inbred mutants. In this study, we challenged this hypothesis by testing whether inbreeding does enhance the suppressive effect of RdDM mutants in the *osd1* background. We report that inbred mutants in *nrpd1, nrpe1* and *drm2* have a successively enhanced ability to suppress the triploid block, but only *nrpd1* can significantly suppress the triploid block in the first generation. Inbreeding of *nrpd1* was connected to a transgenerational loss of non-CG DNA methylation on sites jointly regulated by CHROMOMETHYLASES 2 and 3 (CMT2/3). Our data thus reveal that loss of RdDM function differs in its effect in early and late generations and that Pol IV may act at early stage of triploid block establishment.

## Results

### Inbreeding of RdDM mutants enhanced their ability to rescue the triploid block

The reported difference in the ability to suppress the triploid block between 4x RdDM mutants (Satyaki and Gehring, 2019) and RdDM *osd1* double mutants (Martinez et al., 2018) could be a consequence of different plant growth conditions or due to the different genetic backgrounds. To distinguish between both possibilities, we tested the 4x RdDM mutants for their ability to suppress the triploid block when grown under our conditions. The suppressive effect of 4x *nrpe1* and 4x *drm2* was indeed as strong as the reported effect of *osd1 nrpd1* and 4x *nrpd1* (Martinez et al., 2018; Satyaki and Gehring, 2019), while we only observed a medium suppressive effect for 4x *dcl3* and a weak suppressive effect for 4x *rdr2* (Supplemental Figure 1). Thus, different growth conditions could not completely explain the difference between the results obtained with RdDM mutants in the *osd1* or 4x background. While the effect of *rdr2* was similar in the *osd1* and 4x background (Martinez et al., 2018) and this study), there was a strong discrepancy for the effect of *nrpe1* and *drm2*. One possible explanation for this difference could be inbreeding; while 4x RdDM mutants were generated by colchicine treatment of inbred RdDM mutants, RdDM mutants in the *osd1* background were tested after the first generation of homozygosity. To test this hypothesis, we analyzed whether inbreeding of RdDM mutants in the *osd1* background would change their ability to suppress the triploid block. We found that inbreeding indeed significantly increased the suppressive effect of *osd1 nrpd1*, *osd1 nrpe1*, *osd1 drm2*, but had only a weak effect on *osd1 rdr2* (Figure 1). Interestingly however, only *osd1 nrpd1* could suppress the triploid block in the F2 generation (first generation of *nrpd1* homozygosity), in contrast to the other RdDM pathway mutants that only had an effect in the F3 and later generations.

**Figure 1.**
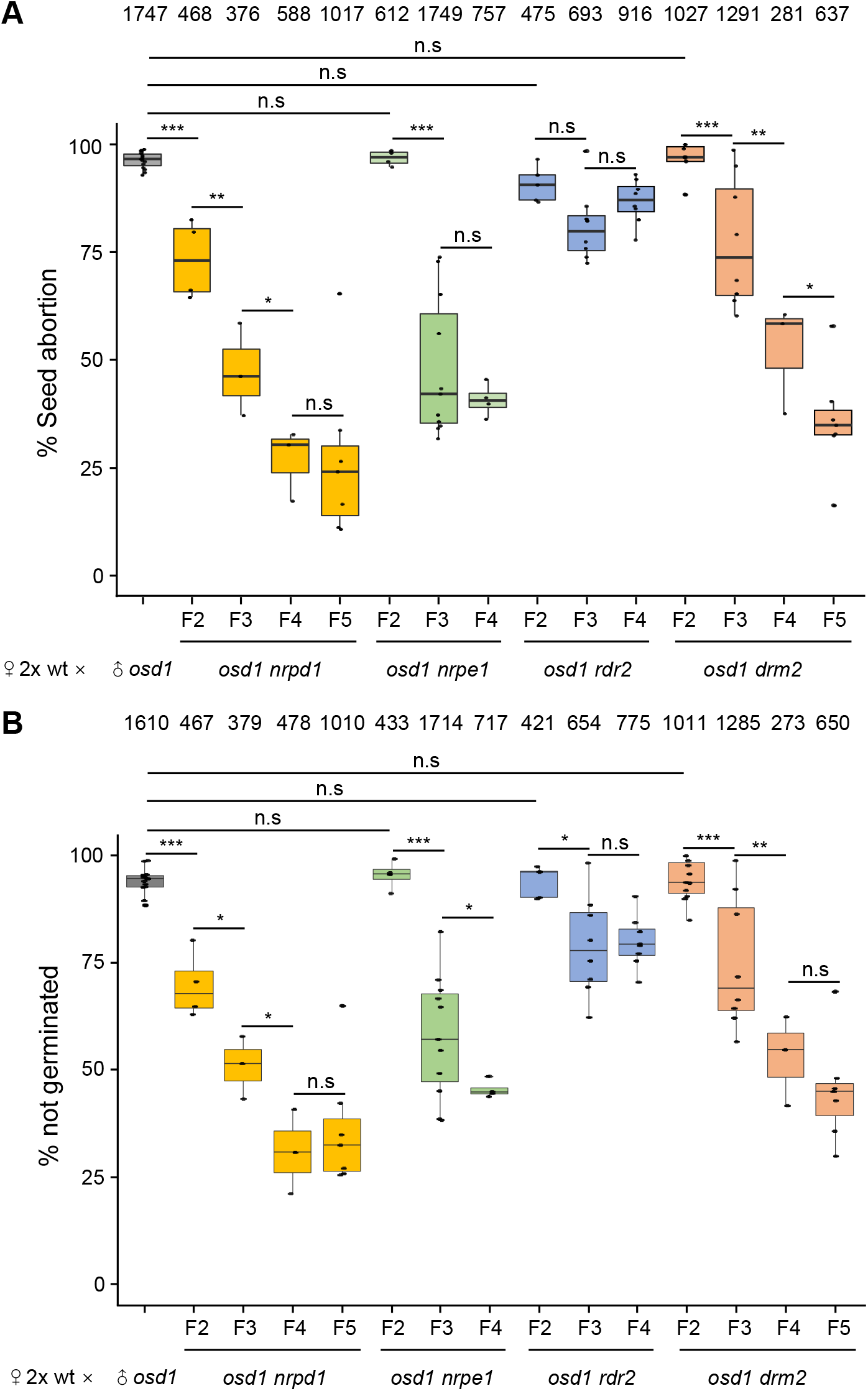
Inbreeding of RdDM *osd1* mutants enhanced their ability to rescue the triploid block. **(A)** Each dot represents the seed abortion rate of 2-4 siliques from a single inflorescence pooled as one cross. Numbers on top represent total seed numbers. **(B)** Each dot represents the percentage of seeds that failed to germinate from each cross. Numbers on top represent total seed numbers. Boxes show medians and the interquartile range, and error bars show the full range. ***, P < 0.001, **, P < 0.01, *, P < 0.05, n.s, not significant (Pairwise t-test).

### Inbreeding of *nrpd1* caused increased loss of DNA methylation

Since the suppressive effect of the triploid block by *osd1 nrpd1*, *osd1 nrpe1* and *osd1 drm2* became stronger with increasing number of generations, we suspected that there is a transgenerational change of DNA methylation from F2 to higher inbred generations of *nrpd1*. We tested this hypothesis by generating bisulfite sequencing data of first generation homozygous *nrpd1* mutants segregating in an F2 population and higher inbred generations of *nrpd1* (more than three times inbred, denoted as Fi) and tested for differences in DNA methylation (Figure 2A, Supplemental Table 1). The RdDM pathway targets cytosines in all sequence contexts, but has its main effects on non-CG methylation (Stroud et al., 2014). We therefore focussed on transgenerational changes in CHG and CHH methylation in F2 and inbred (Fi) *nrpd1* mutants. We identified differentially methylated regions (DMRs) that were hypomethylated in first generation homozygous *nrpd1* (F2) mutants compared to wild type (referred to as DMR1, see Figure. 2A, Supplemental Dataset 1), hypomethylated DMRs between F2 and inbred (Fi) *nrpd1* mutants (referred to as DMRi, Supplemental Dataset 1), and hypomethylated DMRs between *nrpd1* Fi mutants and wild type (referred to as DMRx, Supplemental Dataset 1). As expected, most DMR1 were also detectable in inbred generations (DMRx) (Figures 2A to 2C). However, we found that inbred generations of *nrpd1* mutants gained many DMRs in CHG and CHH sequence contexts (DMRi, Figures 2D and 2E).

**Figure 2.**
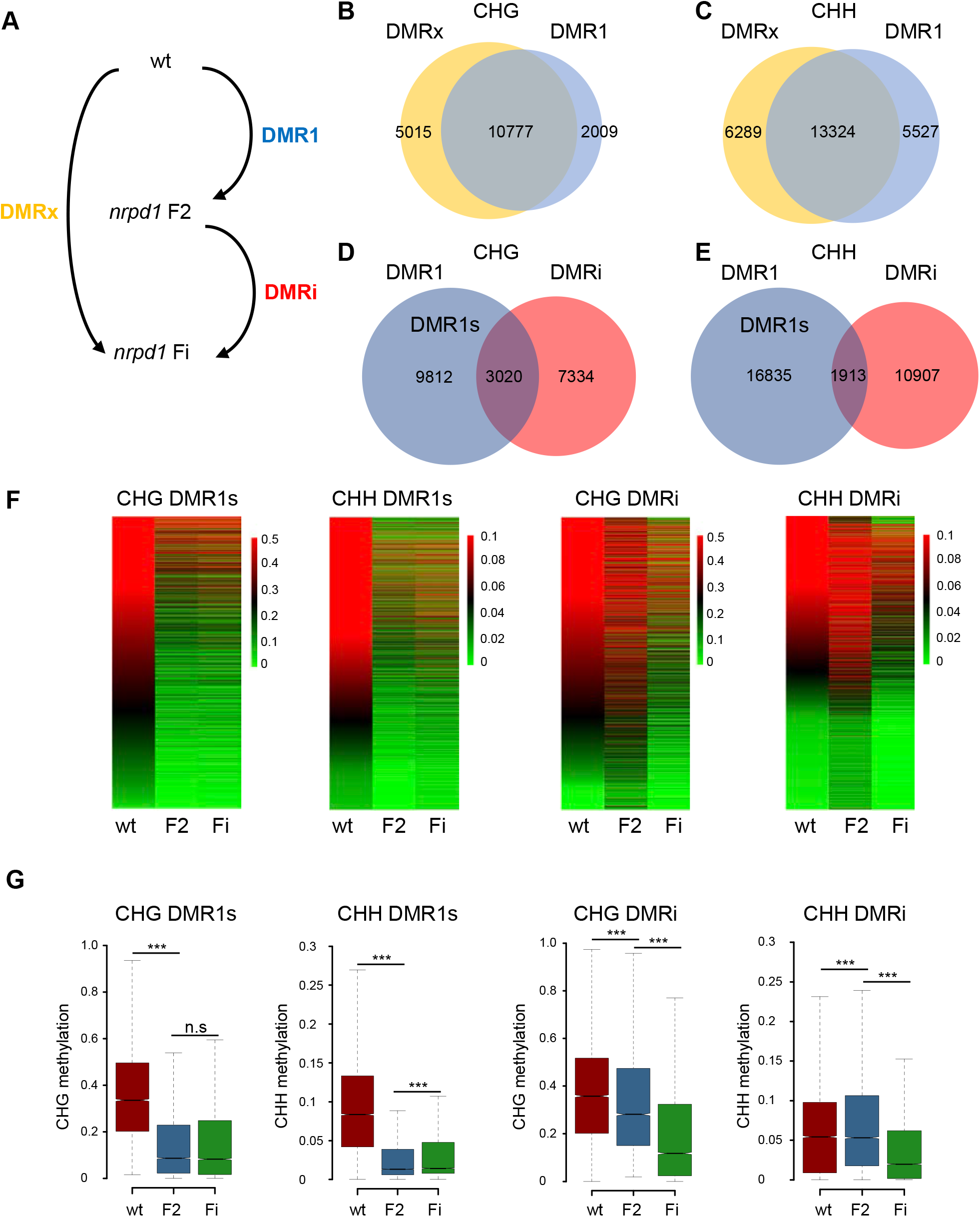
Transgenerational change of non-CG DNA methylation in inbred generations of *nrpd1* mutants. **(A)** Scheme of defining three groups of differentially methylated regions (DMRs). Wild type (wt), first generation homozygous *nrpd1* (F2) and inbred *nrpd1* (Fi). **(B)** and **(C)** Venn diagrams showing the overlap of CHG DMRx and DMR1 **(B)** and CHH DMRx and DMR1 **(C)**. **(D)** and **(E)** Venn diagrams showing the overlap of CHG DMR1 and DMRi **(D)** and CHH DMR1 and DMRi **(E)**. DMR1s refers to DMR1 not overlapping with DMRi. **(F)** Heatmaps of CHG and CHH DNA methylation levels at DMR1s and DMRi loci in wt, F2 and Fi *nrpd1* mutants. **(G)** Boxplots of CHG and CHH DNA methylation levels at DMR1s and DMRi loci in wt, F2 and Fi *nrpd1* mutants. Boxes show medians and the interquartile range, and error bars show the full range excluding outliers. ***, P < 0.001, n.s, not significant (Wilcoxon test).

Those DMR1 regions that were not affected by inbreeding and were thus not overlapping with DMRi regions were defined as DMR1s (Figures 2D and 2E) and compared to DMRi. Visualization of DMR1s and DMRi using heatmaps and boxplots revealed instant loss of DNA methylation in the first generation of *nrpd1* homozygous mutants for DMR1s and gradual loss of DNA methylation upon inbreeding for DMRi (Figures 2F and 2G), consistent with the defining criteria for DMR1s and DMRi.

In wild-type plants, methylation levels in CHG context were slightly higher in DMRi compared to DMR1s, while CHH methylation levels were significantly lower in DMRi compared to DMRs (Figure 3A), suggesting differential activity of the RdDM pathway on both types of DMRs. Nearly half of DMR1s and DMRi associated with TEs, the other half associated with promoter and coding regions (Figure 3B). There were significant differences in the association of both types of DMRs to genic regions; DMRi were more frequently associated with coding regions than DMR1s and conversely, DMR1s were more frequently associated with promoter regions than DMRi (Figure 3B). DMR1s and DMRi were preferentially associated with different TE families; CHG and CHH DMR1s were more frequently associated with helitrons, but depleted in Gypsy and Copia TEs. Conversely, DMRi were more frequently associated with Gypsy TEs, but depleted on helitrons (Figure 3C).

**Figure 3.**
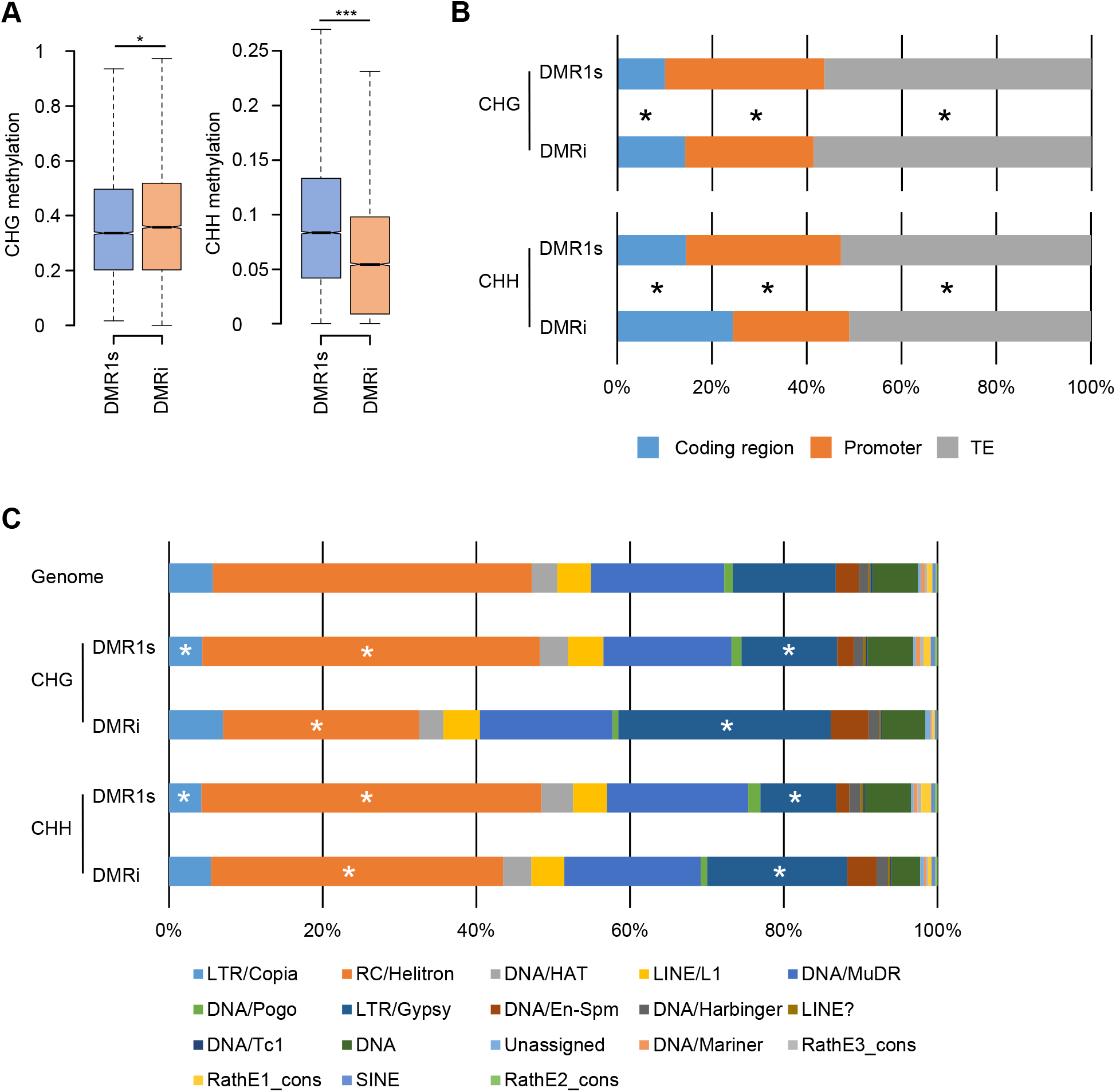
Different DNA methylation levels of DMR1s and DMRi in wild type and their distribution over genomic features. **(A)** Boxplots of CHG and CHH DNA methylation levels at DMR1s and DMRi in wild type. Boxes show medians and the interquartile range, and error bars show the full range excluding outliers. ***, P < 0.001, *, P < 0.05 (Wilcoxon test). **(B)** Distribution of non-CG DMR1s and DMRi over genomic features. Asterisks denote significant differences between DMR1s and DMRi (P < 0.001; Chi square test). **(C)** Percentage of TE families intersected with non-CG DMR1s and DMRi. Asterisks denote significant differences between observed and expected enrichments of TE families (P < 0.001; Chi square test).

### Loss of RdDM differentially affects DMR1s and DMRi

Using published bisulfite data of various mutants in RdDM components and other DNA methylation pathways (Stroud et al., 2013), we tested whether DMR1s and DMRi were differentially affected by loss of different silencing pathways. Indeed, DMR1s and DMRi differed in their response to loss of RdDM pathway mutants; loss of CHG and CHH methylation was significantly stronger in *nrpd1*, *nrpe1*, *rdr2*, and *drm1/2* on DMR1s than on DMRi (Figures 4A and 4B, Supplemental Figures 2A and 2B). This data indicate that DMR1s differ from DMRi in their dependency on RdDM and that methylation at DMRi is redundantly maintained by RdDM and other mechanisms. Previous work revealed that CHG and CHH methylation are partially redundantly regulated by all non-CG methyltransferases, which includes DRM2, CMT2 and CMT3 (Stroud et al., 2014). We therefore analyzed CHG and CHH methylation on DMR1s and DMRi in *cmt2* and *cmt3* mutant backgrounds (Figures 4A and 4B, Supplemental Figures 2A and 2B). Since CMT2 and CMT3 are recruited by H3K9me2 (Jackson et al., 2002; Ebbs and Bender, 2006; Du et al., 2012), we included the H3K9me2 depleted *suvh4/5/6* triple mutant in this analysis. Consistent with the idea that DMRi is redundantly targeted by other DNA methylation pathways, we found that DMRi experienced a significantly stronger loss of CHG methylation in *cmt3* and *suvh4/5/6* mutant backgrounds compared to DMR1s (Figure 4A). Similarly, CHH methylation levels in DMRi were significantly stronger affected by loss of *CMT2* than in DMR1s (Figure 4B). Nevertheless, despite the stronger effect of *suh4/5/6*, *cmt3*, and *cmt2* on DMRi, also DMR1s were significantly affected in those mutants (Supplemental Figure 2). Preferential targeting of DMRi by CMT2 and CMT3 pathways correlated with significantly higher levels of H3K9me2 on DMRi compared to DMR1s (Figures 4C and 4D). Conversely, DMR1s had significantly higher levels of 24-nt sRNAs compared to DMRi (Figures 4C and 4D), correlating with their preferential targeting by the RdDM pathway. Together, we conclude that DMR1s and DMRi are redundantly targeted by RdDM, CMT2 and CMT3 pathways (Figures 4A and 4B, Supplemental Figures 2A and 2B). While loss of RdDM components had a stronger effect on DMR1s, DMRi were more strongly affected by loss of the CMT2/CMT3 pathway, providing a possible explanation for the persistent DNA methylation on DMRi upon initial loss of the RdDM pathway.

**Figure 4.**
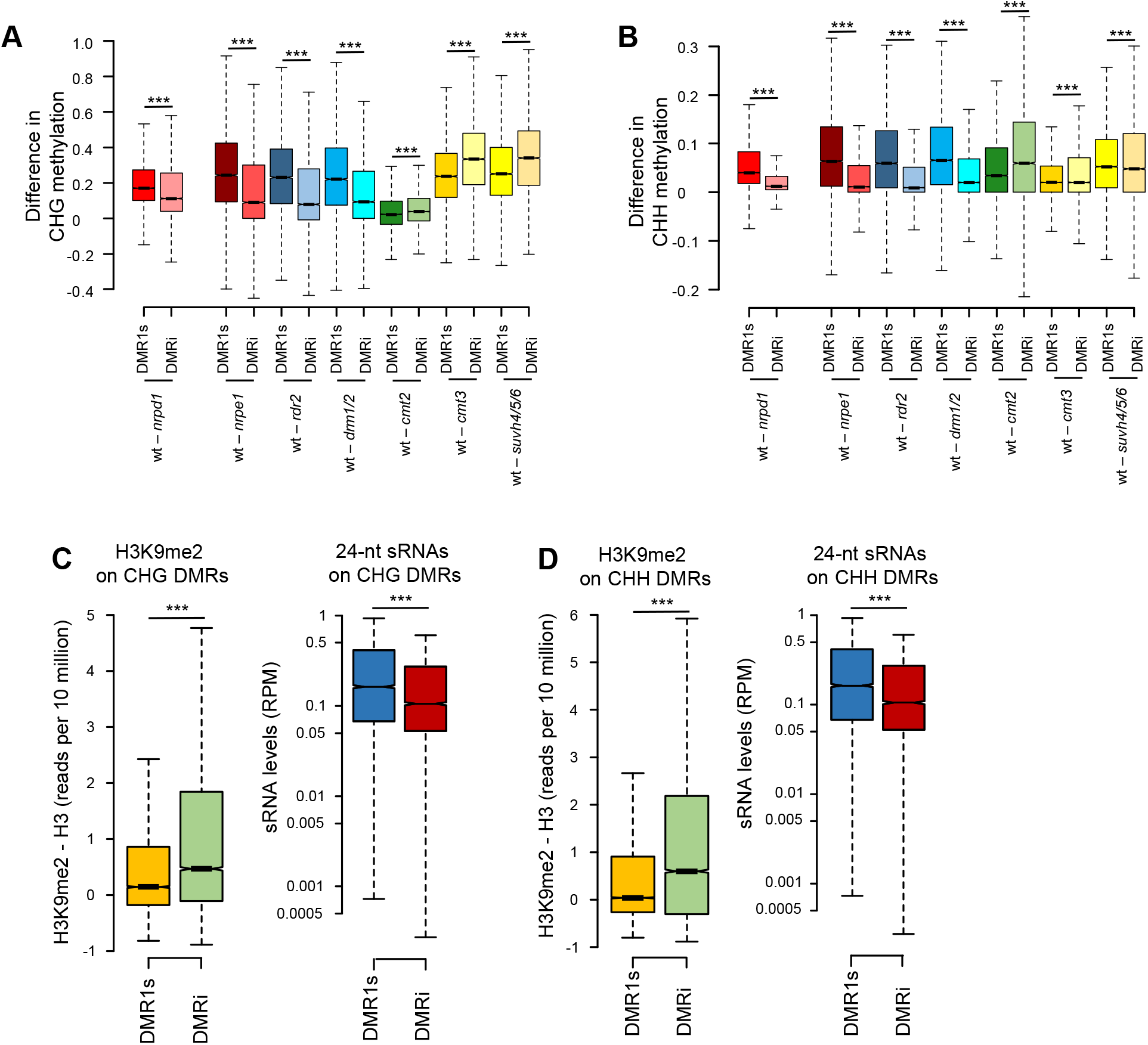
DMR1s and DMRi are differentially affected by loss of RdDM and other DNA methylation pathways. **(A)** Boxplots showing CHG methylation loss in mutants compared to wild type (wt - mutant) on DMR1s and DMRi. **(B)** Boxplots showing CHH methylation loss in mutants compared to wt (wt - mutant) on DMR1s and DMRi. **(C)** Boxplots of H3K9me2 and 24-nt siRNAs levels on CHG DMR1s and DMRi in wt leaves. **(D)** Boxplots of H3K9me2 and 24-nt siRNAs levels on CHH DMR1s and DMRi in wt leaves. Boxes show medians and the interquartile range, and error bars show the full range excluding outliers. ***, P < 0.001 (Wilcoxon test).

### DMRi overlap with deregulated genes in triploid seeds

The *osd1 nrpd1* mutant was able to suppress the triploid block in the first homozygous generation, but the suppressive effect was strongly enhanced by increasing generations of inbreeding (Figure 1). Similarly, *osd1 nrpe1* and *osd1 drm2* gained the ability to suppress the triploid block after successive generations of inbreeding. One possible explanation for this phenomenon is that causal loci affected by NRPD1 lose DNA methylation with increasing generations of inbreeding. To test this hypothesis, we identified genes overlapping DMRi and analyzed their expression in triploid seeds. We found a significant overlap of genes that were upregulated in the endosperm of 3x seeds (log2FC > 1, p < 0.05) and downregulated in the endosperm of 3x *nrpd1* seeds (log2FC < −1, p < 0.05) with those having a CHG DMRi in their vicinity (within 1kb of promoter and coding region) (Figure 5A). There was no significant overlap of deregulated genes with CHH DMRi (Figure 5A). Those genes overlapping with CHG DMRi were strongly enriched for transcription factors, in particular type I AGAMOUS-LIKE (AGL) transcription factors and AUXIN RESPONSE FACTORS (ARFs) (Figure 5B, Supplemental Dataset 2). Among the *AGL* genes was *AGL28*, which encodes a potential interaction partner for PHERES1 (PHE1) and *AGL36*, a close ortholog of *PHE1* that may possibly act redundantly with *PHE1* (Parenicova et al., 2003; Batista et al., 2019b). Loss of PHE1 function can suppress the triploid block (Batista et al., 2019b), implicating that the regulation of AGLs by Pol IV-derived siRNAs is functionally relevant. In support of this notion, we found a significant overlap of genes downregulated in triploid *phe1 phe2* seeds with downregulated genes in 3x *nrpd1* seeds (Figures 5C to 5E, Supplemental Figure 3). The affected *ARFs 12, 13, 14, 20-23* are specifically expressed in the endosperm and belong to a cluster of ARFs located close to the centromere of chromosome 1 (Rademacher et al., 2011). High auxin levels antagonize endosperm cellularization (Batista et al., 2019a) and endosperm cellularization failure is the main cause for triploid seed arrest (Hehenberger et al., 2012). Since ARFs mediate auxin signalling, increased ARF expression in 3x seeds may connect to endosperm cellularization failure, while negative regulation of ARFs in triploid *nrpd1* seeds may lead to restored endosperm cellularization.

**Figure 5.**
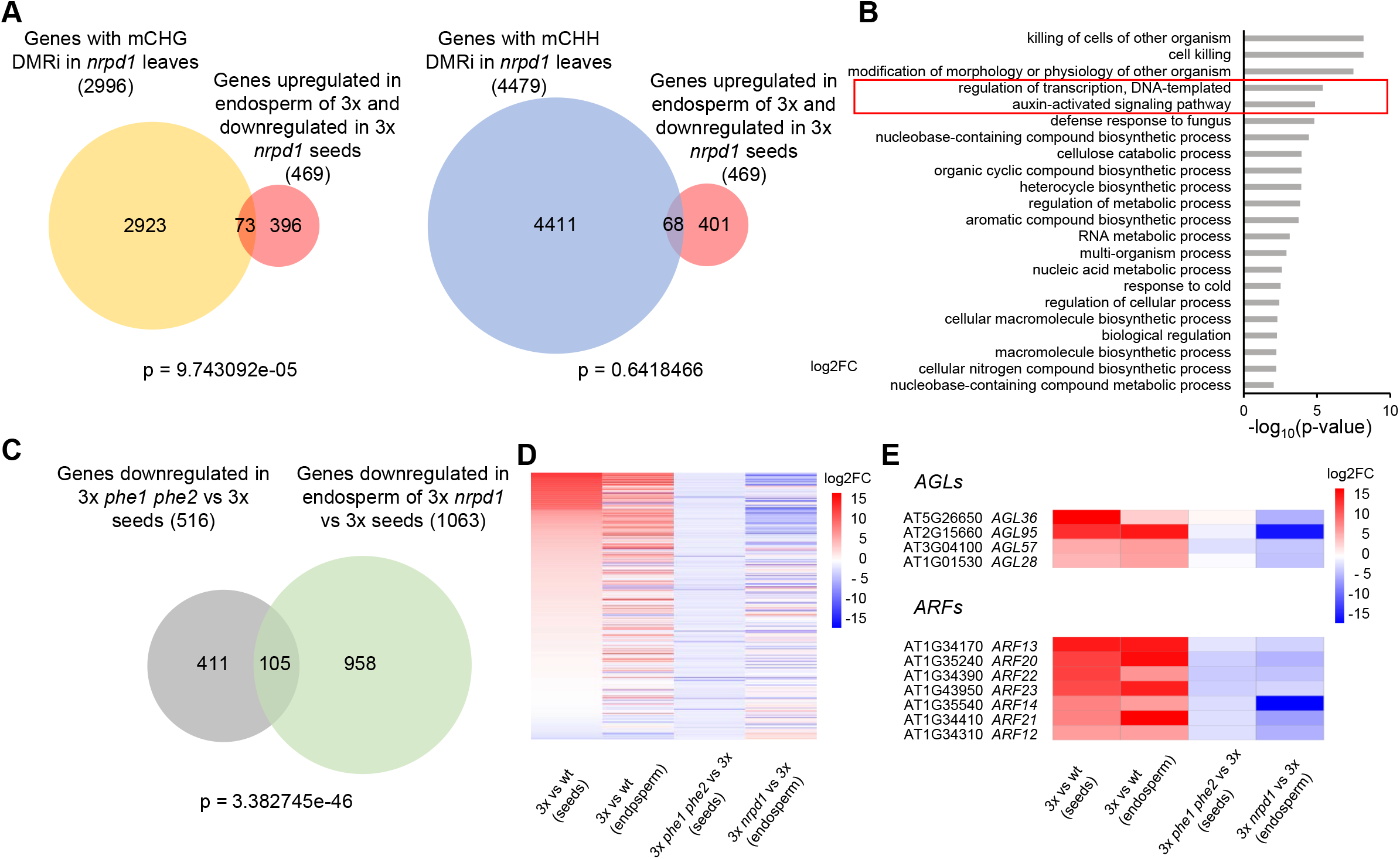
Overlap of DMRi with deregulated genes in triploid (3x) seeds. **(A)** Venn diagrams showing overlap of non-CG DMRi intersected genes and deregulated genes in *Arabidopsis* endosperm of 3x seeds (log2FC > 1, p < 0.05 in 3x vs wt and log2FC < −1, p <0.05 in 3x *nrpd1* vs 3x). **(B)** Enriched gene ontologies (GO) for biological processes (p-value < 0.01) of intersected CHG DMRi overlapping genes and deregulated genes in *Arabidopsis* endosperm of 3x seeds (log2FC > 1, p < 0.05 in 3x vs wt and log2FC < −1, p <0.05 in 3x *nrpd1* vs 3x). **(C)** Venn diagram showing overlap of genes downregulated (log2FC < −1, p <0.05) in 3x *phe1 phe2* seeds vs 3x seeds (Batista et al. 2019) and genes downregulated (log2FC < −1, p <0.05) in endosperm of 3x *nrpd1* vs 3x seeds (Martinez et al. 2018). **(D)** Heatmap showing genes downregulated (log2FC < −1, p <0.05) in 3x *phe1 phe2* seeds vs 3x seeds (Batista et al. 2019) and their expression in endosperm of 3x vs wt seeds (Martinez et al. 2018), 3x *phe1 phe2* vs 3x seeds, and endosperm of 3x *nrpd1* vs 3x seeds (Martinez et al. 2018). **(E)** Heatmap of type I *AGLs* and *ARFs* with CHG DMRi and upregulated (log2FC > 1, p <0.05) in endosperm of 3x vs wt seeds (Martinez et al. 2018), and their expression in 3x vs wt and 3x *phe1 phe2* vs 3x seeds (Batista et al. 2019) and endosperm of 3x *nrpd1* vs 3x seeds.

### Genes marked by DMRi undergo DNA methylation changes in the endosperm of 3x seeds

We tested whether genes overlapping with DMRi would be similarly marked by DMRs in the endosperm of 3x versus 2x seeds. The endosperm of 3x seeds has reduced CHH methylation, which is partly restored upon pollination with *nrpd1* pollen (Martinez et al., 2018; Satyaki and Gehring, 2019). Using previously published data (Satyaki and Gehring, 2019), we identified genes overlapping with hypomethylated DMRs (hypo DMRs) in the endosperm of 3x versus 2x and hypermethylated DMRs (hyper DMRs) in 3x *nrpd1* versus 3x seeds. A significant number of genes that overlapped with CHG and CHH DMRi in *nrpd1* leaves also had CHG and CHH hypo DMRs in the endosperm of 3x seeds (Figures 6A). Importantly, among those genes were *AGL28*, *AGL36* and *ARFs 12-14, 20-23* (Figure 6B, Supplemental Figure 4, Supplemental Dataset 3). A significant number of genes marked by CHH hypoDMRs in the endosperm of 3x seeds gained CHH methylation in the endosperm of 3x *nrpd1* seeds (Figure 6C), among those *ARFs 12, 14, 15, 20-23*.

**Figure 6.**
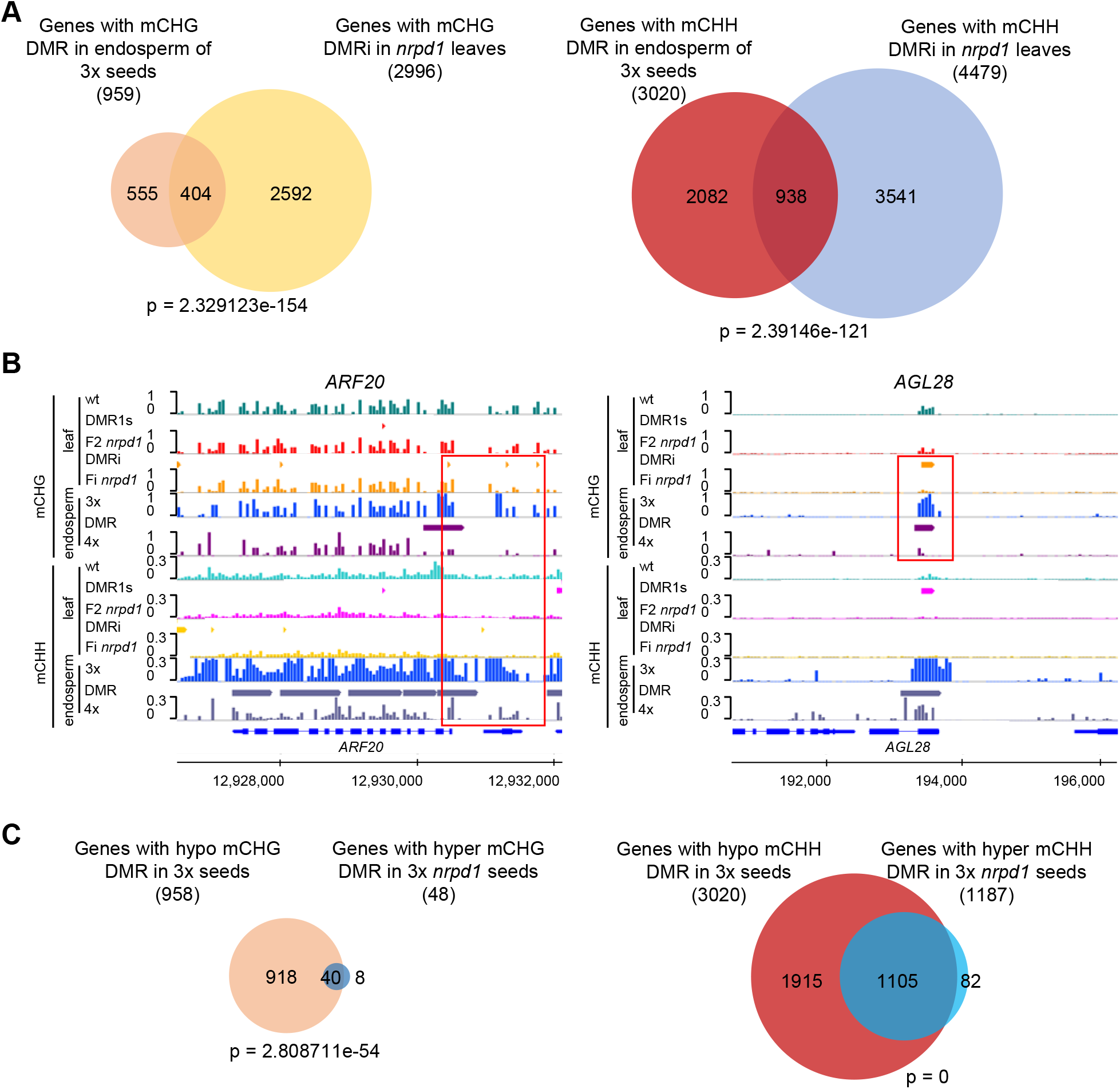
DMRi associated with genes losing DNA methylation in the endosperm of 3x seeds. **(A)** Venn diagrams showing overlap of genes with hypo non-CG DMRs in the endosperm of 3x seeds (Satyaki and Gehring 2019) and genes with non-CG DMRi in *nrpd1* leaves. **(B)** Representative *ARF* and *AGL* losing non-CG methylation during generations in *nrpd1* leaves and endosperm of 3x seeds (Satyaki and Gehring 2019). Representative regions are highlighted by a red box. DMRs are marked by bars. **(C)** Venn diagrams showing overlap of genes with hypo non-CG DMRs in the endosperm of 3x versus 2x seeds and hyper DMRs in endosperm of 3x *nrpd1* versus 3x seeds (Satyaki and Gehring 2019).

Together, our data reveal that inbreeding of *nrpd1*, *nrpe1* and *drm2* in the *osd1* background triggered increased suppression of the triploid block. Inbreeding of *nrpd1* caused increasing loss of CHG and CHH methylation at defined loci, providing a possible explanation for the enhanced suppressive effect of RdDM mutants over generations. Nevertheless, *nrpd1* was able to suppress the triploid block in the F2 generation, differing from *nrpe1* and *drm2*, suggesting that Pol IV acts at an early stage of triploid block establishment independent of its role in RdDM.

## Discussion

In this study, we report that mutations in the RdDM components *NRPD1*, *NRPE1* and *DRM2* triggered a successively enhanced suppression of the triploid block over generations of inbreeding. Inbreeding of *nrpd1* caused an increasing loss of CHH and CHG methylation over generations, suggesting that the generation-dependent suppression of the triploid block connects to generation-dependent loss of DNA methylation. Loci showing increasing loss of CHG and CHH methylation over generations (DMRi) were marked by higher levels of H3K9me2 and were more strongly affected in *cmt2* and *cmt3* mutants than loci that lost DNA methylation in the first homozygous generation (DMR1s) of *nrpd1*. This suggests that DMRi loci are partly redundantly targeted by RdDM, CMT2 and CMT3 pathways and that upon loss of RdDM the efficiency of CMT2 and CMT3 to maintain methylation on those loci decreases over generations. The *suvh4/5/6* triple mutant has a strong suppressive effect on the triploid block (Jiang et al., 2017), supporting a possible redundancy of CMT3 and CMT2 pathways on functionally relevant loci. Cooperation of all non-CG methyltransferases to regulate CHG and CHH methylation was previously demonstrated in *Arabidopsis* (Stroud et al., 2014), adding support to this notion.

Interestingly, we found that only *nrpd1* had a suppressive effect on the triploid block in the first homozygous generation, differing from *nrpe1* and *drm2* that required one additional generation to have an effect. During pollen development, Pol IV generates an abundant class of 21/22-nt sRNAs (referred to as epigenetically activated RNAs, easiRNAs) that we previously proposed to act as the dosage-dependent signal inducing the triploid block (Borges et al., 2018; Martinez et al., 2018). This view has been challenged by data revealing that 4x mutants in aforementioned RdDM components suppress the triploid block (Satyaki and Gehring, 2019). However, as shown in this and our previous study, mutants defective in core components of the RdDM pathway, *rdr2*, *nrpe1* and *drm2*, do not have a suppressive effect in the first homozygous generation (Martinez et al., 2018) and this study. This renders DNA methylation as the sole transmissible signal for genome dosage rather unlikely. We thus propose that Pol IV acts very early in the pathway leading to the triploid block, possibly by generating sRNAs that establish a transmissible signal for genome dosage (Figure 7). Pol IV is recruited to sites that are redundantly targeted by RdDM, CMT2 and CMT3 pathways. This scenario could explain why loss of Pol IV function leads to a suppressive effect in the first homozygous generation, while loss of RdDM pathway components have a suppressive effect only in later generations. If Pol IV target sites remain methylated in the first generation of RdDM mutants (as shown for DMRi), they would still be able to recruit Pol IV, which would generate the signal and induce the triploid block (Figure 7). Once DNA methylation on Pol IV target sites is lost, which will happen after inbreeding of RdDM mutants, failure of Pol IV recruitment will abolish the signal and the triploid block is not triggered. We therefore propose that relevant DMRs establishing the triploid rescue are only erased after several rounds of inbreeding of RdDM mutants, thus corresponding to DMRi.

**Figure 7.**
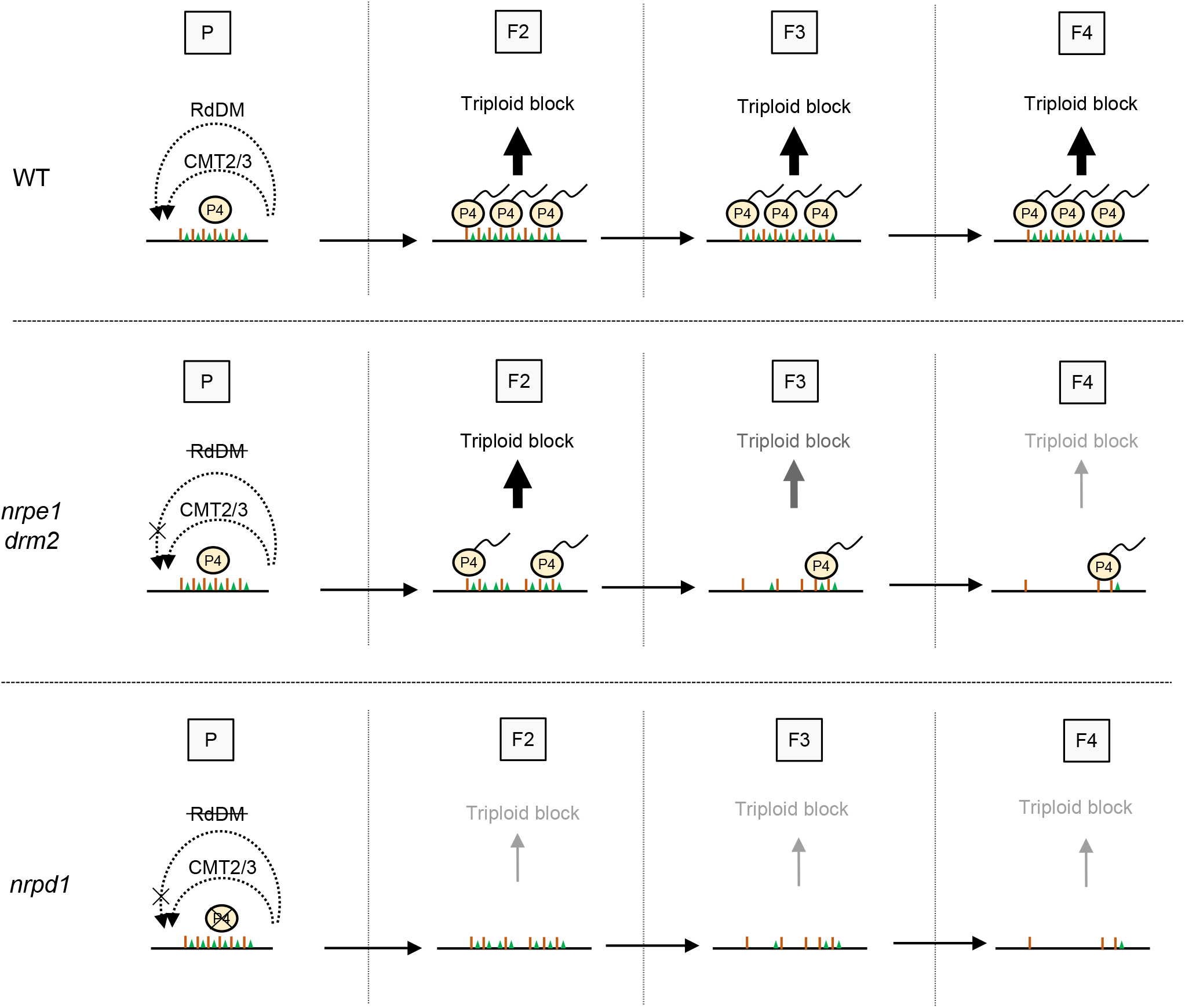
Hypothetical model describing the link between transgenerational loss of CHH methylation in *nrpd1* and the triploid block. Causal loci for the triploid block are targeted by the RdDM and the CMT2/CMT3 pathways (referred to as DMRi in the manuscript), leading to stable DNA methylation. Pol IV (P4) is recruited to those loci and produces the signal(s) able to induce the triploid block. In RdDM mutants like *nrpe1* or *drm2*, DNA methylation at DMRi gradually decreases over generations. This leads to a reduction of Pol IV recruitment, reducing the strength of the triploid block in a transgenerational way. In *nrpd1*, the signal is not produced and the rescue of the triploid block can be observed in the first generation. However, as for the other RdDM mutants, the DNA methylation will decrease over generations. P4: Pol IV, brown bars represent DNA methylation, green triangles represent H3K9me2.

Interestingly, we found DMRi to overlap with genes that are potentially relevant for establishing the triploid block, like AGLs and ARFs (Batista et al., 2019a; Batista et al., 2019b). Previous work from our group revealed that the AGL PHE1 is a central regulator of imprinted genes and loss of *PHE1* causes strong suppression of the triploid block (Batista et al., 2019b). We found a significant overlap of genes being negatively regulated upon loss of PHE1/PHE2 and NRPD1 function, suggesting a possible connection. AGLs were previously shown to be regulated by maternal 24-nt sRNAs (Lu et al., 2012; Kirkbride et al., 2019), including *AGL28* and *AGL36* that we found to be marked by DMRi and lose DNA methylation in the endosperm of triploid seeds. AGL28 is a potential interaction partner for PHE1, while AGL36 is in the same clade as PHE1 and both AGLs may act redundantly (Parenicova et al., 2003). All three AGLs were highly upregulated in 3x seeds and suppressed by *nrpd1*, correlating with the overlap of deregulated genes in 3x *phe1 phe2* and *nrpd1* seeds. Whether increased dosage of easiRNAs negatively interferes with RdDM as previously proposed, remains to be further tested, but the strong overlap of loci targeted by DMRi and hypoDMRs in the endosperm of triploid seeds suggest a possible connection.

In summary, in this study we reveal that inbreeding of mutants impaired in RdDM components successively enhanced their ability to suppress the triploid block. Only loss of Pol IV function had a suppressive effect in the first homozygous generation; while other RdDM mutants required at least two generations of inbreeding to cause an effect. Our data thus reveal that loss of RdDM function differs in its effect in early and late generations and that the initial signal of genome dosage is tightly coupled to Pol IV function, but not the canonical RdDM pathway.

## Methods

### Plant Growth and Material

*Arabidopsis* mutants *nrpd1-3* (SALK_128428) (Onodera et al., 2005), *nrpe1-12* (SALK_033852) (Pontier et al., 2005), *rdr2-2* (SALK_059661) (Herr et al., 2005), and *drm2-2* (SALK_150863) (Chan et al., 2006) were obtained from Nottingham *Arabidopsis* Stock Centre (NASC). The *osd1-1* mutant (d’Erfurth et al., 2009) was kindly provided by Raphael Mercier. The tetraploid mutants *nrpd1-4* (SALK_083051), *nrpe1-11* (SALK_029919), *rdr2-1* (SAIL_1277_H08), *drm2-2*, and *dcl3-1* (SALK_005512) were kindly shared by Mary Gehring (Satyaki and Gehring, 2019). The Col-0 accession was used as the wild type for all experiments. Primers used for genotyping all mutants are listed in Supplemental table 2. *Arabidopsis* seeds were surface-sterilized in 5% commercial bleach and 0.01% Tween 20 for 10 min, followed by three times washes in sterile distilled, deionized water. Seeds were sown on half-strength Murashige and Skoog medium (0.43% [w/v] Murashige and Skoog salts, 0.8% [w/v] bacto agar, 0.19% [w/v] MES hydrate, and 1% [w/v] Suc). After stratification (2 days at 4°C), the plates were transferred to a growth chamber (16h of light/8h of dark, 110 μmol photons m^−2^ s^−1^, 21°C, 70% humidity). Ten-day old seedlings were transferred to soil and grown in a growth chamber under a 16-h-light/8-h-dark cycle with light intensity of 150 μmol photons m^−2^ s^−1^, at 21°C and 70% humidity.

For crosses, diploid Col-0 wild-type buds were emasculated 2 days before pollination with indicated pollen donors.

### Bisulfite sequencing

*Arabidopsis* three-week old aerial parts were pooled from three plants as one replicate and ground with liquid nitrogen into fine powder and then used for isolation of genomic DNA using the MagJET Plant Genomic DNA Kit (K2761). Duplicates were generated from each sample. Libraries were prepared with the Accel-NGS Methyl-Seq DNA Library Kit from Illumina (Cat No. 30096, Swift) and the sequencing was performed at Novogene (Hongkong, China) on a NovaSeq 6000 platform in 150-bp paired-end mode.

### Chromatin Immunoprecipitation (ChIP) followed by sequencing

Cross-linked *Arabidopsis* leaves (100 mg) from each sample were ground in liquid nitrogen into fine powder and used for further experiments as previously described (Moreno-Romero et al., 2016). Biological triplicates were generated from each sample. Libraries were generated using 1.5 ng of starting material using the Ovation Ultralow Library System (NuGEN, San Carlos, USA) and the sequencing was performed at Novogene (Hongkong, China) on a HiSeqX in 150-bp paired-end mode.

Anti-histone H3 (Sigma, #H9289) and anti-H3K9me2 (Diagenode, #pAb-060-050) antibodies were used in this study.

### Bioinformatic analysis

For DNA methylation analysis, 150-bp paired-end reads were trimmed by removing the first 5 bases from the 5’ end and the last 20 bases from the 3’ end. Reads were mapped to the *Arabidopsis* TAIR10 in paired-end mode (--score_min L,0,−0.6) genome using Bismark (Krueger and Andrews, 2011). Duplicated reads were eliminated and methylation levels for each condition were calculated by averaging the replicates. Differentially methylated regions (DMRs) in CG, CHG and CHH contexts were calculated only for hypomethylation in order wild type > F2 *nrpd1* (DMR1), F2 *nrpd1* > Fi *nrpd1* (DMRi), and wild type > Fi *nrpd1* (DMRx) in 50 bp bins, considering as the fractional methylation threshold the bins with differences below the 1st decile. The bins with p-value < 0.01 (Fisher’s exact test) were considered as significant. DMRs located within 300 bp from each other were merged to yield the final list of DMRs. Genes (gene-body plus 1kb upstream) overlapping with indicated DMRs were obtained using intersect feature of bedtools v2(Quinlan and Hall, 2010). Reads of ChIP-seq samples passing a quality control were mapped to the Arabidopsis (TAIR10) genome using Bowtie (Langmead, 2010) in single-end mode, allowing for up to two mismatches. Mapped reads were deduplicated and extended to the estimated average length of the genomic fragments (270 bp). Coverage was estimated and normalized to 10 million reads. H3K9me2 ChIP signals were normalized by subtracting their coverage with H3 ChIP data at every single position in the genome.

For small RNA analysis, the resulting 18-30-bp-long sRNAs reads after removing adapters were mapped to the *Arabidopsis* TAIR10 genome. After removing reads mapping to chloroplast and mitochondria and to structural noncoding RNAs (tRNAs, snRNAs, rRNAs, or snoRNAs), the resulting mapped reads from both replicates were pooled together, sorted in 21/22-nt and 24-nt categories and remapped to the same reference masked genome mentioned above using ShorStack (–mismatches 0–mmap f) (Johnson et al., 2016) in order to improve the localization of sRNAs mapping to defined DMR loci. The alignments were normalized by converting coverage values to RPM values.

PLAZA 4.0 dicots (Van Bel et al., 2017) was used to identify enriched Gene Ontology (GO) terms. GO terms of biological functions with p-value < 0.01 were further loaded on REVIGO (Supek et al., 2011) to remove the redundant terms. The charts were generated based on −log10 (p-values).

Publicly available datasets used in this study are as follows: methylation data in *Arabidopsis* RdDM mutants was from GSE39901 (Stroud et al., 2013), small RNA data in *Arabidopsis* wild-type and *nrpd1* leaves was from GSE116067 (Tan et al., 2018), RNA-seq data of *Arabidopsis osd1 nrpd1* endosperm was from GSE84122 (Martinez et al., 2018), RNA-seq of *Arabidopsis osd1 phe1 phe2* seeds was from GSE129744 (Batista et al., 2019b), and DMRs and methylation data in the endosperm of *Arabidopsis* wt and *nrpd1* triploid seeds were from GSE126929 (Satyaki and Gehring, 2019).

## Data availability

The sequencing data generated in this study are available in the Gene Expression Omnibus under accession number GSE156597. Supplemental Table 1 summarizes all sequencing data generated in this study.

## Acknowledgments

This research was supported by the Swedish Research Councils VR and Formas (grants 2016-00961 and 2017-04119 to C.K.), the Knut and Alice Wallenberg Foundation (grant 2018-0206 to C.K.), and the Göran Gustafsson Foundation for Research in Natural Sciences and Medicine (to C.K.).

## Author Contributions

Z.W. and C.K. performed the experimental design; Z.W. and C.W. performed experiments; N.B. advised on experimental work; Z.W., C.K., L.S., and J.S-G. analysed data, Z.W., N.B., J.S.-G., and C.K. wrote the article; all authors read and commented on the article.

## Supplemental Figure legends

**Supplemental Figure 1.**
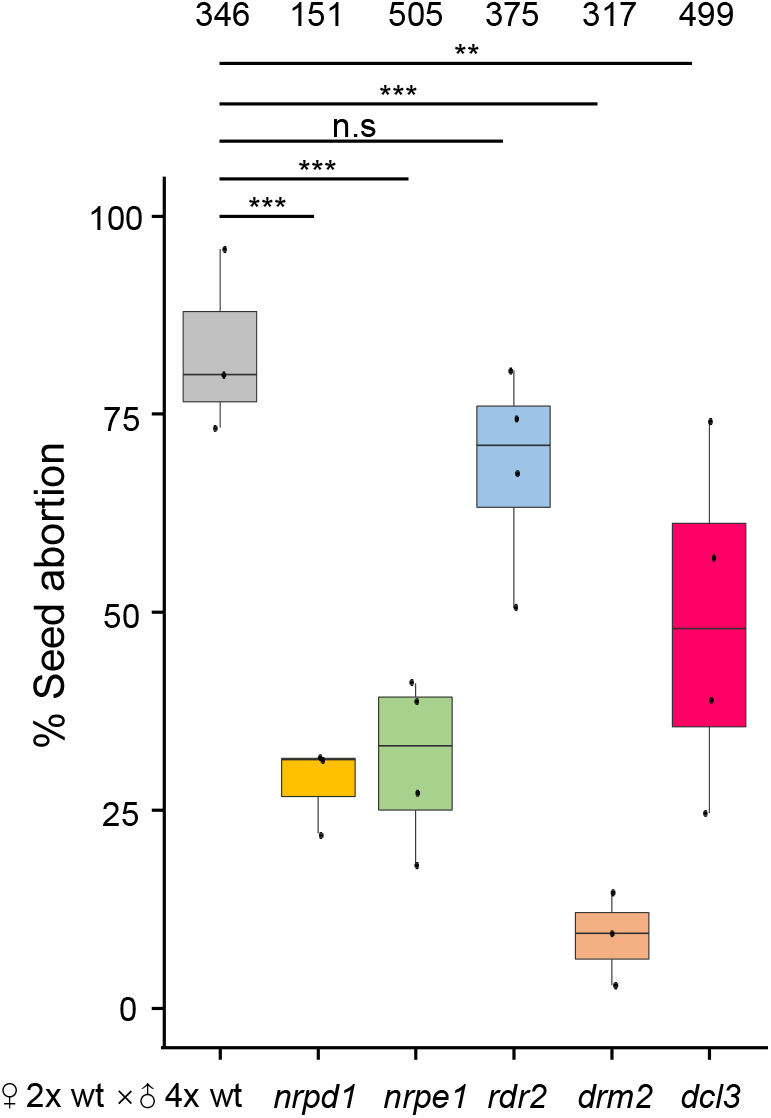
Rescue of 3x seed abortion using 4x RdDM mutants as pollen donor. Each dot represents the seed abortion rate of 2-4 siliques from a single inflorescence pooled as one cross. Boxes show medians and the interquartile range, and error bars show the full range. Numbers on top represent total seed numbers. ***, P < 0.001, **, P < 0.01, n.s, not significant (Pairwise t-test).

**Supplemental Figure 2.**
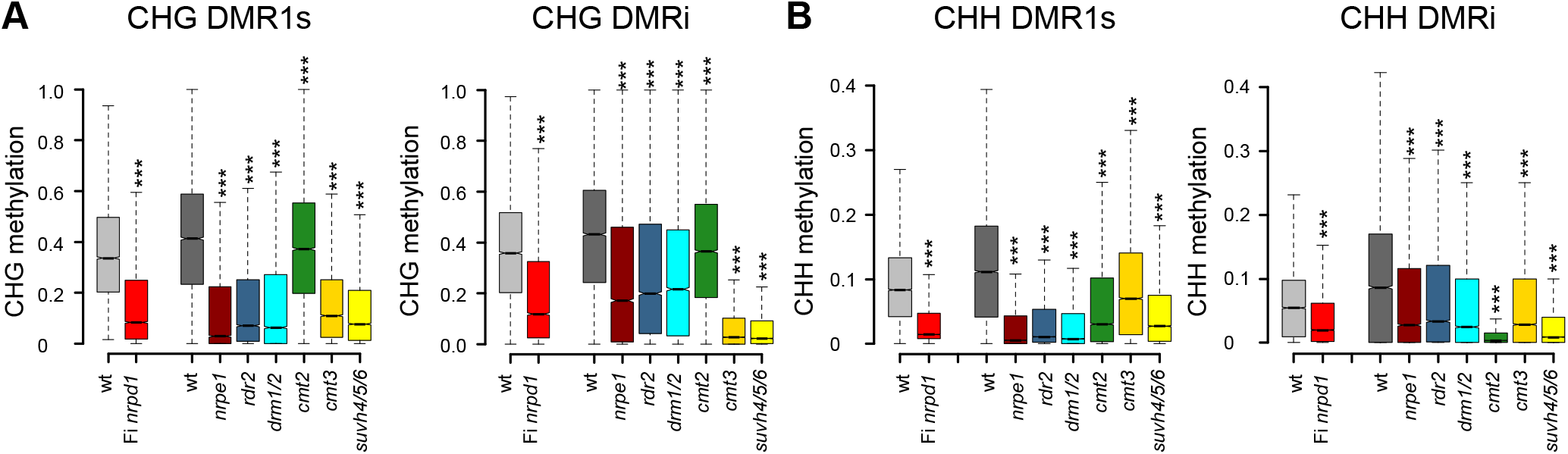
Non-CG methylation on DMR1s and DMRi in mutants of RdDM and other DNA methylation pathways. **(A)** Boxplots of CHG methylation levels on DMR1s and DMRi in wild type (wt) and Fi *nrpd1* (data generated in this study) and DNA methylation mutants (data generated in Stroud et al., 2013). **(B)** Boxplots of CHH methylation levels on DMR1s and DMRi in wt and Fi *nrpd1* (data generated in this study) and DNA methylation mutants (data generated in Stroud et al., 2013). Boxes show medians and the interquartile range, and error bars show the full range excluding outliers. ***, P < 0.001 (Wilcoxon test). The statistical tests were performed between indicated mutants and wt.

**Supplemental Figure 3.**
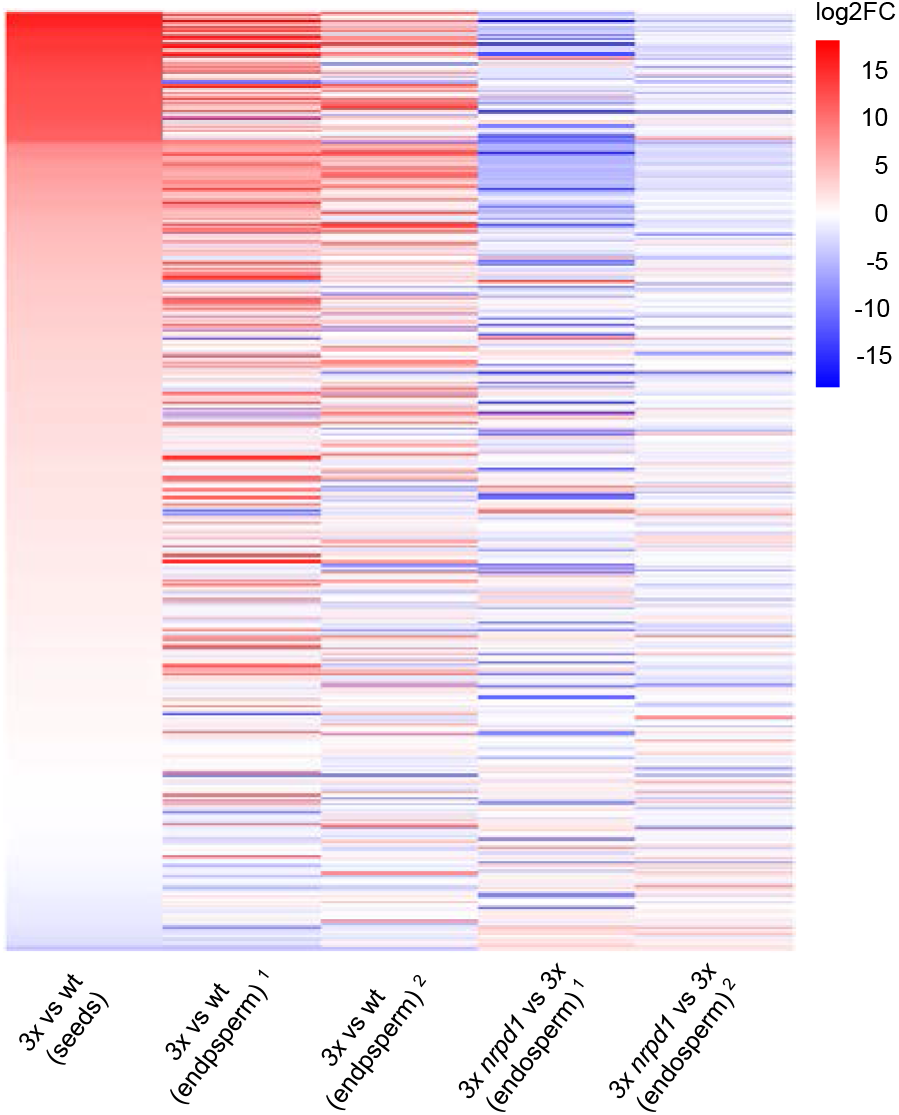
Similar expression pattern of deregulated genes in 3x seeds and the endosperm of 3x seeds. Heatmap showing gene expression of downregulated genes (log2FC < −1, p < 0.05) in 3x *phe1 phe2* vs 3x seeds (Batista et al. 2019) and their expression in 3x vs wt seeds (Batista et al. 2019) and two sources of endosperm of 3x vs wt seeds and endosperm of 3x *nrpd1* vs 3x seeds. Dataset 1 (Martinez et al. 2018) and dataset 2 (Satyaki and Gehring 2019).

**Supplemental Figure 4.**
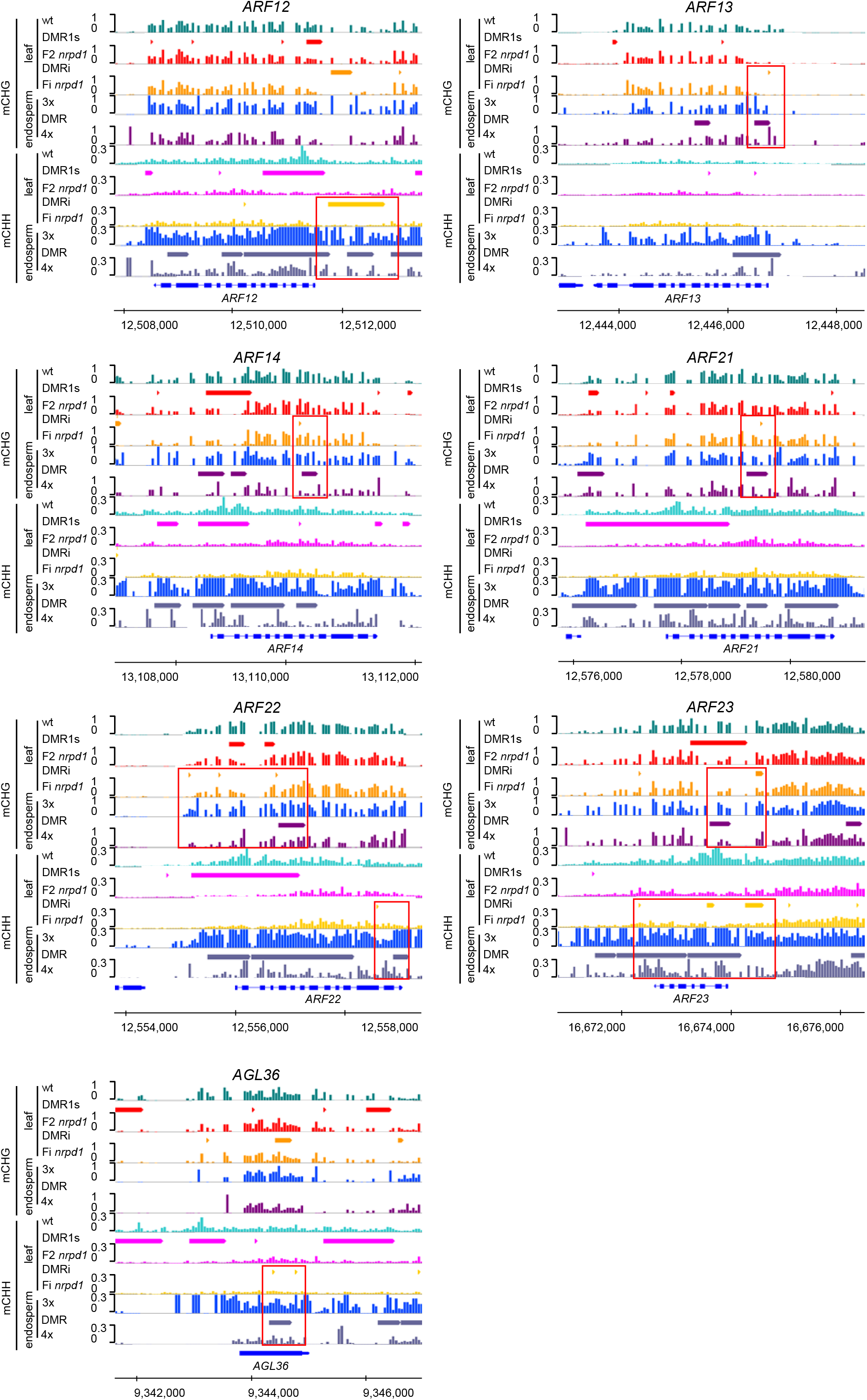
Screenshots of selected *ARFs* and *AGLs* losing non-CG methylation in flanking or coding regions during generations in *nrpd1* leaves and endosperm of 3x seeds (Satyaki and Gehring 2019). Bars on top represent DMRs. Red boxes highlight regions with DMRi.

## Supplemental datasets

**Supplemental Dataset 1**. Differentially methylated regions (DMRs) that were hypomethylated in first generation homozygous *nrpd1* (F2) mutants compared to wild type (DMR1), hypomethylated DMRs between F2 and inbred (Fi) *nrpd1* mutants (DMRi), and hypomethylated DMRs between *nrpd1* Fi mutants and wild type (DMRx).

**Supplemental Dataset 2**. Genes intersected with CHG DMRi overlapping with genes upregulated in the endosperm of 3x vs wt and downregulated in 3x *nrpd1* vs 3x seeds.

**Supplemental Dataset 3**. Genes intersected with non-CG DMRi overlapping with genes with non-CG hypo DMRs in endosperm of 3x seeds.

---

**Supplemental table 1.**
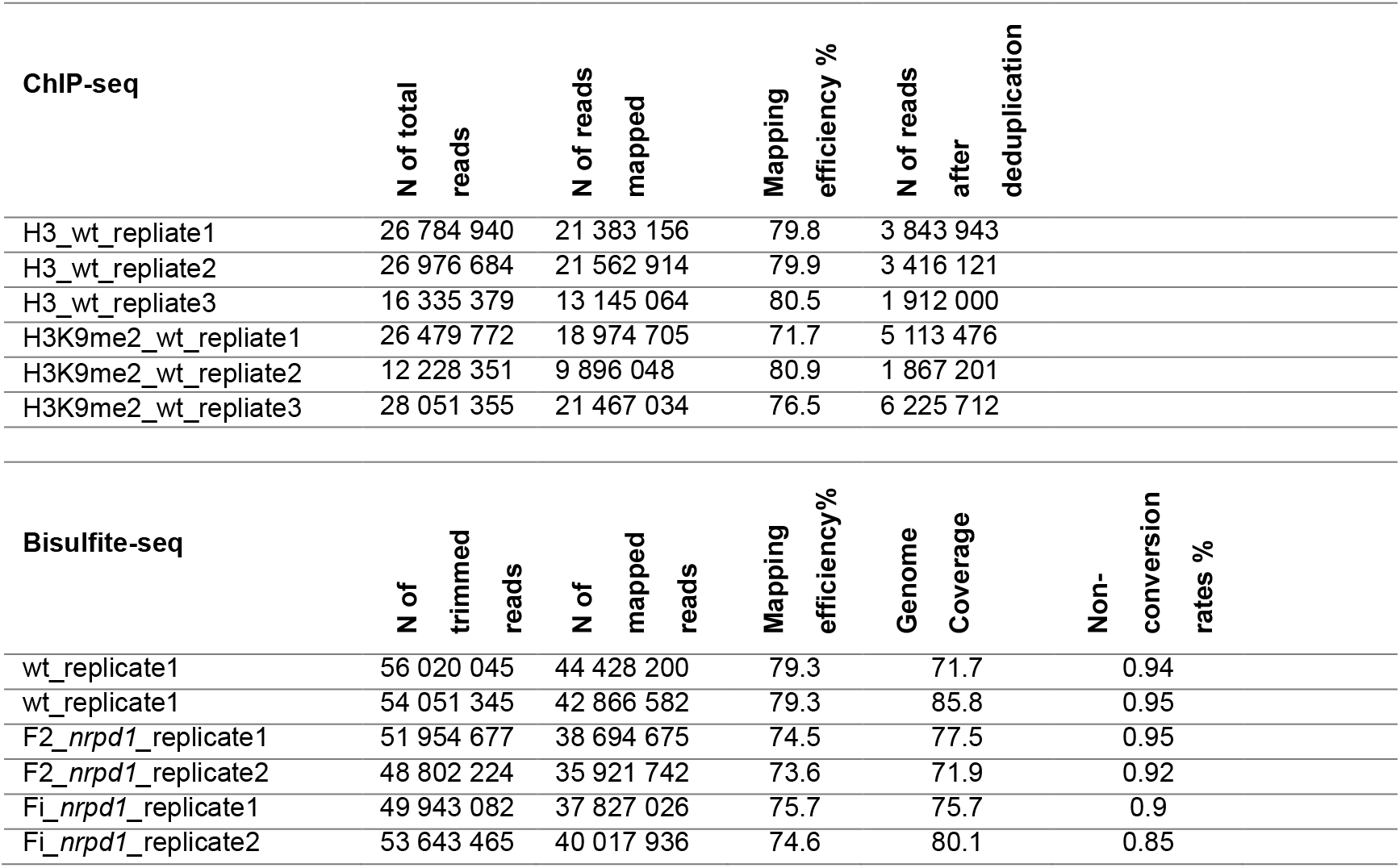
Quality of sequencing samples. Table shows details of the sequenced samples generated in this study. Replicates are biological replicates.

**Supplemental table 2.**
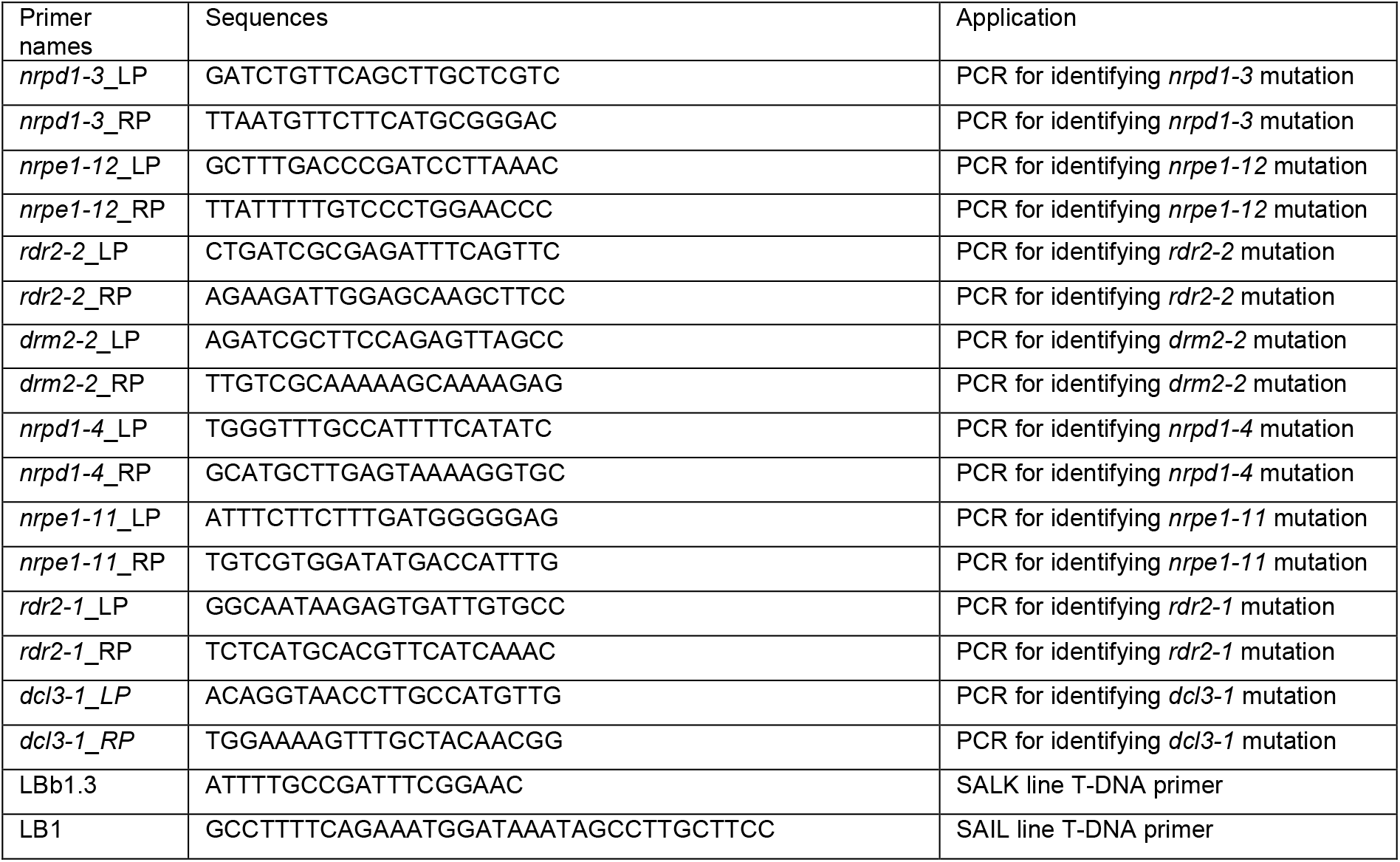
Primer list.

